# Anxiety-like but not despair-like behaviors are further aggravated by chronic mild stress in the early stages of APP_swe_/PS1dE9 transgenic mice

**DOI:** 10.1101/202283

**Authors:** Jun-Ying Gao, Ying Chen, Dong-Yuan Su, Charles Marshall, Ming Xiao

## Abstract

Early Alzheimer’s disease (AD) and depression share many symptoms, thus it is very difficult to initially distinguish one from the other. Therefore, characterizing the shared and different biological changes between the two disorders will be helpful in making an early diagnosis and planning treatment. In the present study, 8-week-old APP_swe_/PS1dE9 transgenic mice received chronic mild stress (CMS) for 8 weeks followed by a series of behavioral, biochemical and pathological analyses. APPswe/PS1dE9 mice demonstrated despair- and anxiety-like behaviors, and reduced sociability, accompanied by high levels of soluble beta-amyloid, glial activation, neuroinflammation and brain derived neurotrophic factor signaling disturbance in the hippocampus. Notably, APPswe/PS1dE9 mice exposure to CMS further aggravated anxiety-like behaviors rather than hopelessness and sociability deficits, accompanied with more severe neuroinflammation, and low serum corticosterone increased to the normal level. These results may help to understand the pathogenic mechanism of psychiatric symptoms associated with early AD.

## INTRODUCTION

With an increasingly aging population, the incidence of Alzheimer’s disease (AD) escalates every year, becoming one of the most serious threats to the health of elderly all over the world. During the early stages of AD, a considerable portion of patients are misdiagnosed with depression, due to the overlap in symptoms, such as loss of interest, social withdrawal, sleeping disturbance and impaired concentration (Kessing et al., 2012; Kaiser et al., 2014; Park et al., 2015). Moreover, antidepressants, including selective serotonin reuptake inhibitors, are often not as effective at treating depression symptoms with AD as they are at treating depression alone (Khundakar and Thomas, 2015). Therefore, characterizing the shared and different biological changes between the two disorders will be helpful in making an early diagnosis and planning treatment.

Excessive accumulation of beta-amyloid (Aβ), caused by an imbalance between its production and clearance, is the chief cause of AD occurrence (Huang and Mucke, 2012; Naj et al., 2017). Meanwhile, stressful life events are closely associated with the onset of depression, although the exact mechanisms for this psychological disease are not clear (vanPraag, 2005; Lang and Borgwardt, 2013; Mahar et al., 2014). Despite different pathogeneses, some similar pathological changes are observed between AD and depression, including neuroinflammation and impaired neurotrophic support, which might contribute to depression-related symptoms in the early stages of AD (Murray et al., 2014; Novais and Starkstein, 2015; Quaranta et al., 2015; Ménard et al., 2016; Mushtaq et al., 2016). However, there is a lack of the direct experimental evidence to support this hypothesis.

APPswe/PS1dE9 mice that over express the Swedish mutation of amyloid precursor protein (APP) together with presenilin1 (PS1) deleted in exon 9 display progressive age-related increases in brain parenchymal beta-amyloid β (Aβ) load and cognitive dysfunction (Trinchese et al., 2004; Reiserer et al., 2007). This AD mouse model also exhibits non-cognitive behavioral abnormalities, including anxiety, hyperactivity, and social behavior deficits (Pietropaolo et al., 2012; Bilkei-Gorzo, 2014; Janus et al., 2015; Olesen et al., 2016). However, there is no systematic analysis of mental behaviors in APP/PS1 mice before the onset of cognitive impairment, warranting investigation into the exact pathophysiology.

The chronic mild stress (CMS) is one of most widely used models for replicating major depression. By repeated exposure to an array of varying, unpredictable, and mild stressors for 5 to 8 weeks, mice will develop depression-like and anxiety-like behaviors, accompanied with hyperactivity of the hypothalamic-pituitary adrenal (HPA) axis (Willner, 2016; Nguyen et al., 2017). However, it remains unclear whether CMS aggravates mental abnormalities in APP/PS1 mice before amyloidosis onset.

To address these issues, 8-week of CMS was performed on 16-week old APPswe/PS1dE9 mice, following by analyses of depression/anxiety-related behaviors, levels of serum corticosterone (CORT), and Aβ load, neuroinflammation and brain derived neurotrophic factor (BDNF) signaling pathway in the hippocampus. Our results characterized psychological phenotype in the early stages of APPswe/PS1dE9 mice, accompanied by neuroinflammation and BDNF dysregulation, rather than overactivation of the HPA axis. We also demonstrated that CMS further aggravates anxiety-like behaviors, but not despair-like behaviors of APP_Swe_/PS1_dE9_ mice, with further elevated proinflammatory cytokine interleukin-6 (IL-6) and tumor necrosis factor-α (TNF-α) in the hippocampus.

## RESULTS

### CMS reduced body weight gain in both WT and APPswe/PS1dE9 mice

WT-Control mice and APPswe/PS1dE9-Control mice gained weight following an 8-week feeding in the intact condition. In contrast, both WT-CMS and APPswe/PS1dE9-CMS mice showed loss of weight (treatment, F_1,52_ = 64.029, P = 0.001; genotype, F_1,52_ = 0.051, P = 0.823; interaction, F_1,52_ = 0.125, P = 0.725) (Fig. 1).

**Fig. 1.**
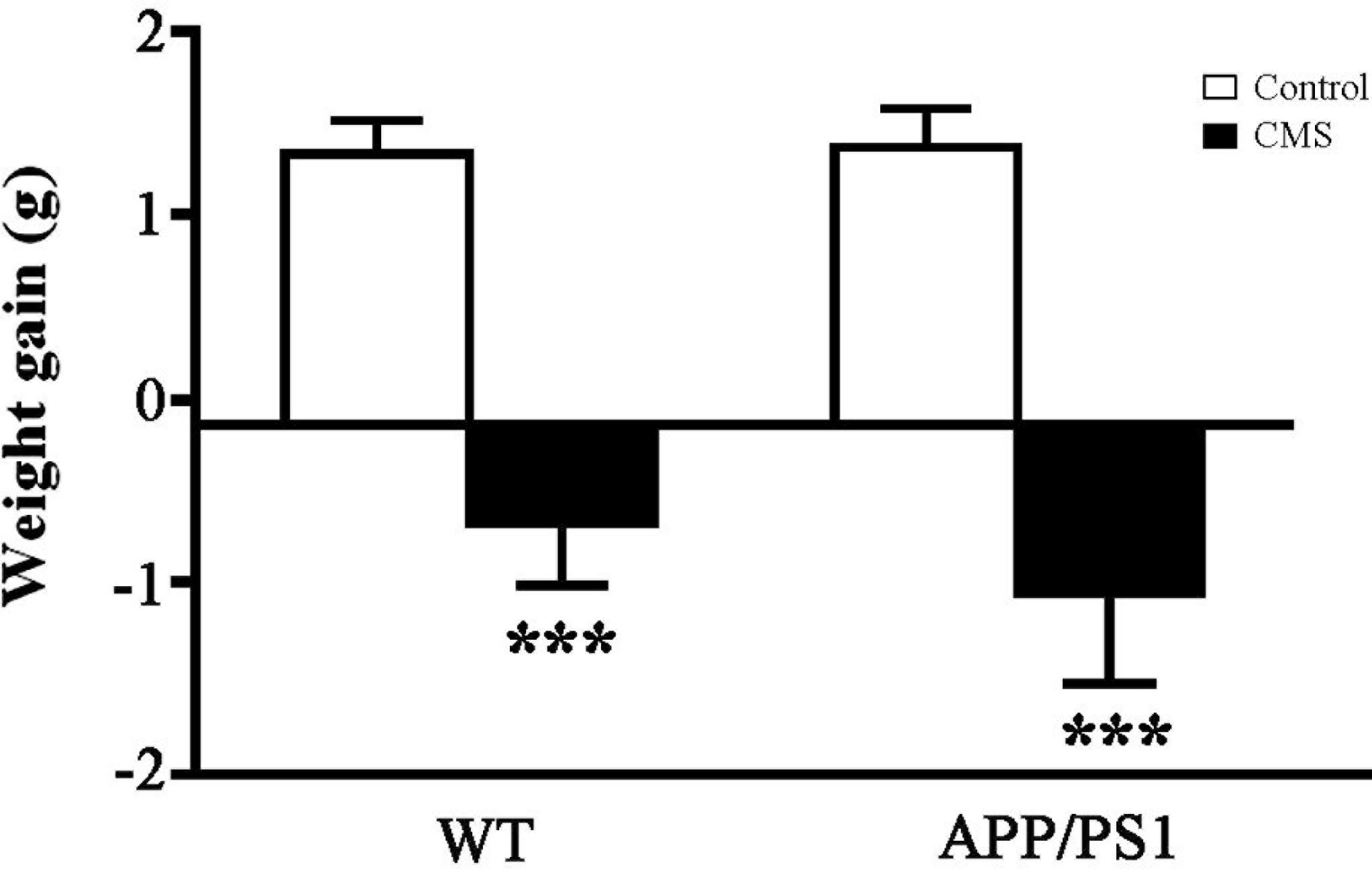
Analyses of weight changes in WT and APPswe/PS1dE9 (APP/PS1) mice by CMS for 8 weeks. Weight change in the eighth week, compared to a baseline starting weight. Data represent means ± SEM. Two-way ANOVA followed by post-hoc multiple comparison test. *P < 0.05; ***P < 0.001, compared to Control. n = 14 in each group.

### CMS exposure did not exacerbate despair-like behaviors of APPswe/PS1dE9 mice

The sucrose preference test was used to investigate anhedonia of each group. CMS significantly reduced the ratio of sucrose solution to total liquid consumed in WT mice, but had no significant effect on APPswe/PS1dE9 mice (treatment, F_1,52_ = 6.069, P = 0.017; genotype, F_1,52_ = 0.431, P = 0.514; interaction, F_1,52_ = 4.795, P = 0.033) (Fig. 2A). These data suggested that anhedonia-like behavior was not observed in APPswe/PS1dE9 mice with or without CMS exposure.

**Fig. 2.**
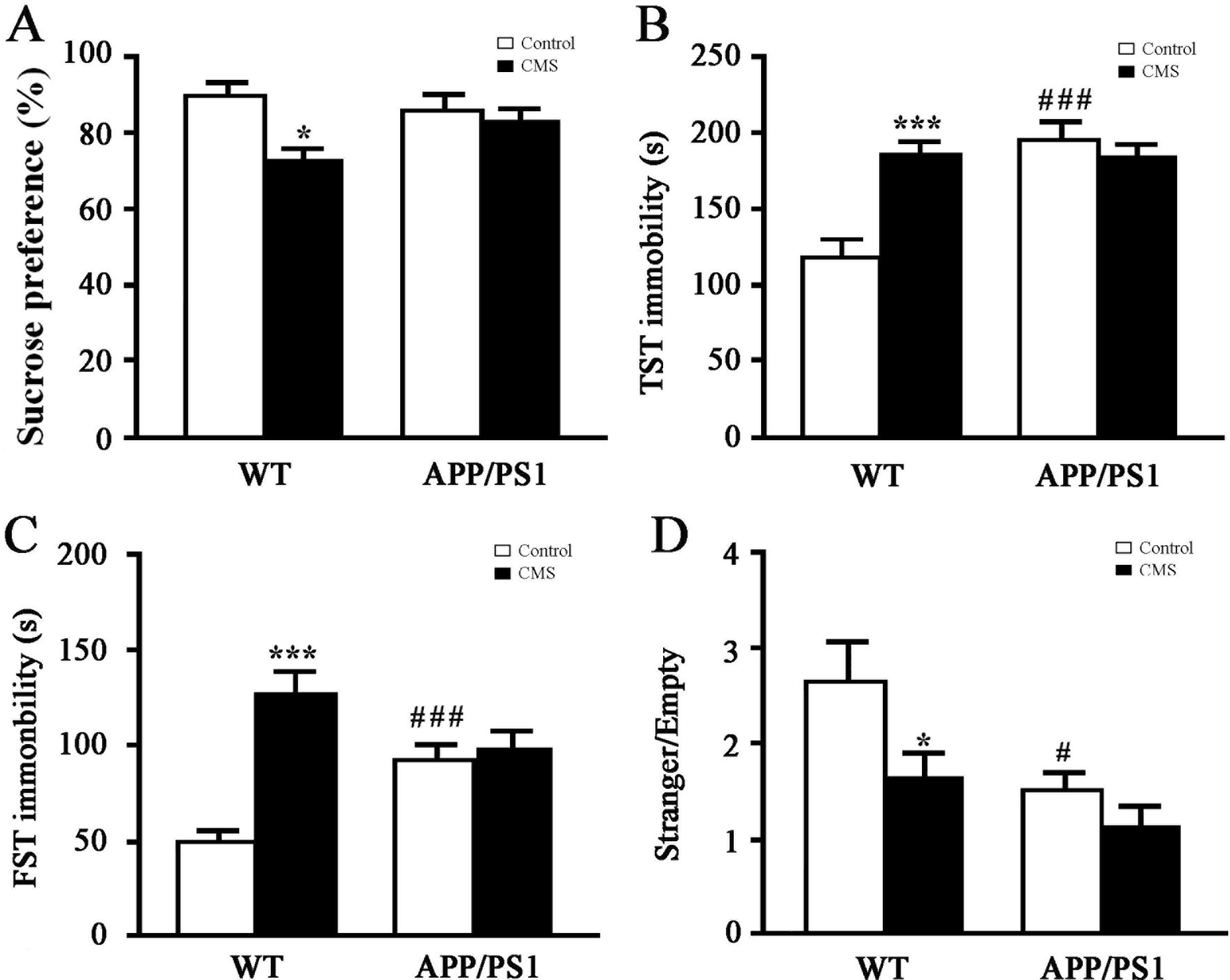
Analysis of depression-like behaviors of WT and APPswe/PS1dE9 (APP/PS1) mice by CMS for 8 weeks. (A) Sucrose preference test. (B) Tail suspension test (TST). (C) Forced swimming test (FST). (D) Social approach test. Data represent means ± SEM. Two-way ANOVA followed by post-hoc multiple comparison test. *P < 0.05; ***P < 0.001, compared to control; #P <0.05; ###P < 0.001, compared to WT. n = 14 in each group.

We then performed the tail suspension test and forced swim test to valuate behavioral despair of mice. Both WT-CMS mice and APPswe/PS1dE9-control mice demonstrated higher immobility time of tail suspension test than WT-Control mice (both P < 0.001) (treatment, F_1,52_ = 8.025, P = 0.007; genotype, F_1,52_ = 11.074, P = 0.002; interaction, F_1,52_ = 15.602, P < 0.001). However, the duration of immobility was not significantly different between APPswe/PS1dE9-Control mice and APPswe/PS1dE9-CMS mice (P > 0.05) (Fig. 2B).

Similarly, in the forced swim test, the immobility time of APPswe/PS1dE9-Control mice was longer than that of WT-Control mice (P < 0.05). CMS significantly increased the immobility time of WT mice, but not APPswe/PS1dE9 mice (treatment, F_1,52_ = 22.511, P < 0.001; genotype, F_1,52_ = 0.643, P = 0.427; interaction, F_1,52_ = 18.569, P < 0.001) (Fig. 2C).

### Anomalies in social behaviors of APPswe/PS1dE9 mice were not affected by CMS

Both patients suffering from depression and patients with early stage AD often exhibit social withdraw (Ballard and Corbett, 2010; Davidson et al., 2016). Anomalies in social behaviors are also present in CMS mouse model and APPswe/PS1dE9 mice (Hsiao et al., 2014; Erburu et al., 2015). In the present study, we further examined their effects on the sociability of mice. Both APPswe/PS1dE9-control mice and WT-CMS mice were less willing to engage in social approach than WT-Control mice, reflected by a low ratio between time spent with an unfamiliar mouse and time in the empty enclosure (CMS, F_1,52_ = 8.379, P = 0.006; genotype, F_1,52_ = 10.931, P = 0.002; interaction, F_1,52_ = 1.740, P = 0.193, respectively). However, CMS did not exacerbate deficits of social interaction in APPswe/PS1dE9 mice (P > 0.05) (Fig. 2D). Together, these results indicated that APPswe/PS1dE9 mice already exhibited psychological abnormalities at 4-month old, which was not exacerbated by CMS exposure.

### CMS increased anxiety-like behaviors of APPswe/PS1dE9 mice

Depression is often complicated by symptoms of anxiety (Melton et al., 2016; Kircanski et al., 2017). Likewise, increased anxiety levels are also observed in CMS mice or rats (Boufleur et al., 2013; Strekalova et al., 2004; Chen et al.,2016; Koprdova et al., 2016), but effects of CMS on anxiety-like behaviors of APPswe/PS1dE9 mice remain unclear. Thus, we selected the open field test and elevated plus maze test to evaluate the anxiety state of mice in each group. CMS significantly decreased the percentage of stay time within the central area (treatment, F_1,52_ = 39.831, P < 0.001; genotype, F_1,52_ = 16.574, P <0.001; interaction, F_1,52_ = 2.040, P = 0.160), as well as the number of entries into the central area in the both genotype mice (treatment, F_1,52_ = 17.689, P < 0.001; genotype, F_1,52_ = 0.349, P = 0.557; interaction, F_1,52_ = 0.349, P = 0.557). APPswe/PS1dE9 mice spent less time in the central area than WT-Control mice (P < 0.05; Fig. 3A), but the entering number was not different between the groups (P > 0.05; Fig. 3B). In addition, CMS reduced the total movement distance in WT mice, but not in APPswe/PS1dE9 mice (treatment, F_1,52_ = 10.068, P < 0.001; genotype, F_1,52_ = 32.098, P < 0.001; interaction, F_1,52_ = 6.817, P = 0.012). APPswe/PS1dE9-Control mice exhibited hyperactivity with long movement distance, compared with WT-Control mice (P < 0.05) (Fig. 3C).

**Fig. 3.**
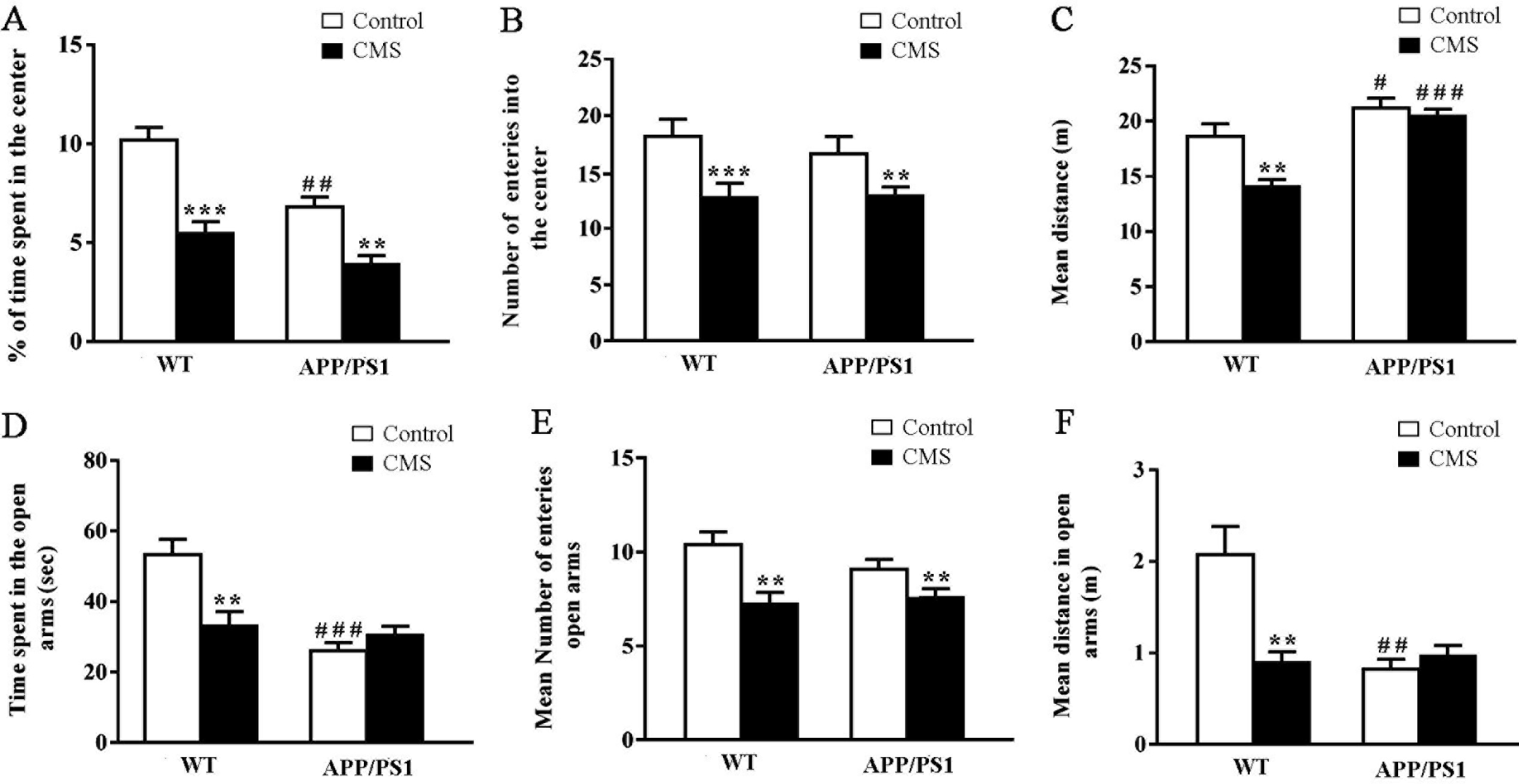
Analyses of anxiety-like behaviors of WT and APPswe/PS1dE9 (APP/PS1) mice by CMS for 8 weeks. (A-C) The open field test. (A) The percent of time spent in the central area. (B) The number of entries into the central area. (C) The movement distance. (D-F) The elevated plus-maze. (D) The percent of time spent in the open arm. (E) The number of entries in the open arm. (F) The total movement distance. Two-way ANOVA followed by post-hoc multiple comparison test. **P < 0.01; ***P < 0.001, compared to Control; #P < 0.05; ##P < 0.01; ###P < 0.001, compared to WT. n = 14 in each group.

In the elevated maze test, CMS reduced the time stayed within the open arm of WT-Control mice, but not APPswe/PS1dE9 mice (treatment, F_1,52_ = 17.576, P < 0.001; genotype, interaction, F_1,52_ = 4.451, P = 0.04; F_1,52_ = 12.411, P = 0.001). APPswe/PS1dE9-Control mice spent less time in the open arm than WT-Control mice (P < 0.01) (Fig. 3D). CMS caused a significant decrease in the entering number in the open arm in the both genotype mice (treatment, F_1,52_ = 16.129, P < 0.001; genotype, F_1,52_ = 0.822, P = 0.369; interaction, F_1,52_ = 2.031, P = 0.161) (Fig. 3E). CMS only reduced the distance of movement in the open arm in WT mice (treatment, F_1,52_ = 8.071, P = 0.007; genotype, F_1,52_ = 2.602, P = 0.113; interaction, F_1,52_ = 5.430, P = 0.024) (Fig. 3F).

Together, CMS exposure exacerbated anxiety-levels of APPswe/PS1dE9 mice, as revealed by decreases in percentage of time spent in and number of entries into the central area, as well as number of entries into the open arm.

### CMS did not impair short-term spatial memory of APPswe/PS1dE9 mice

The Y-maze testing was conducted to examine the short-term spatial memory of mice. Time spent in the novel arm was not affected by CMS (F_1,52_ = 0.860, P = 0.358), genotype (F_1,52_= 0.089, P = 0.911) or their interaction (F_1,52_ = 2.676, P = 0.108) (Fig. 4A). CMS significantly decreased the number of entries into the novel arm of WT mice (P < 0.01), but not APPswe/PS1dE9 mice (treatment, F_1,52_ = 15.975, P < 0.001; genotype, F_1,52_ = 6.218, P = 0.016; interaction, F_1,52_ = 1.637, P = 0.207) (Fig. 4B). APPswe/PS1dE9-CMS mice had higher entering numbers than WT-CMS mice (P < 0.01), suggesting that CMS partially impaired the short-term spatial memory of WT mice rather than APPswe/PS1dE9 mice. In addition, APPswe/PS1dE9 mice showed hyperactivity, reflected by long distance traveled in the novel arm (treatment, F_1,52_ = 1.915, P = 0.173; genotype, F_1,52_ = 32.528, P < 0.0001; interaction, F_1,52_ = 1.559, P = 0.218) (Fig. 4C).

**Fig. 4.**
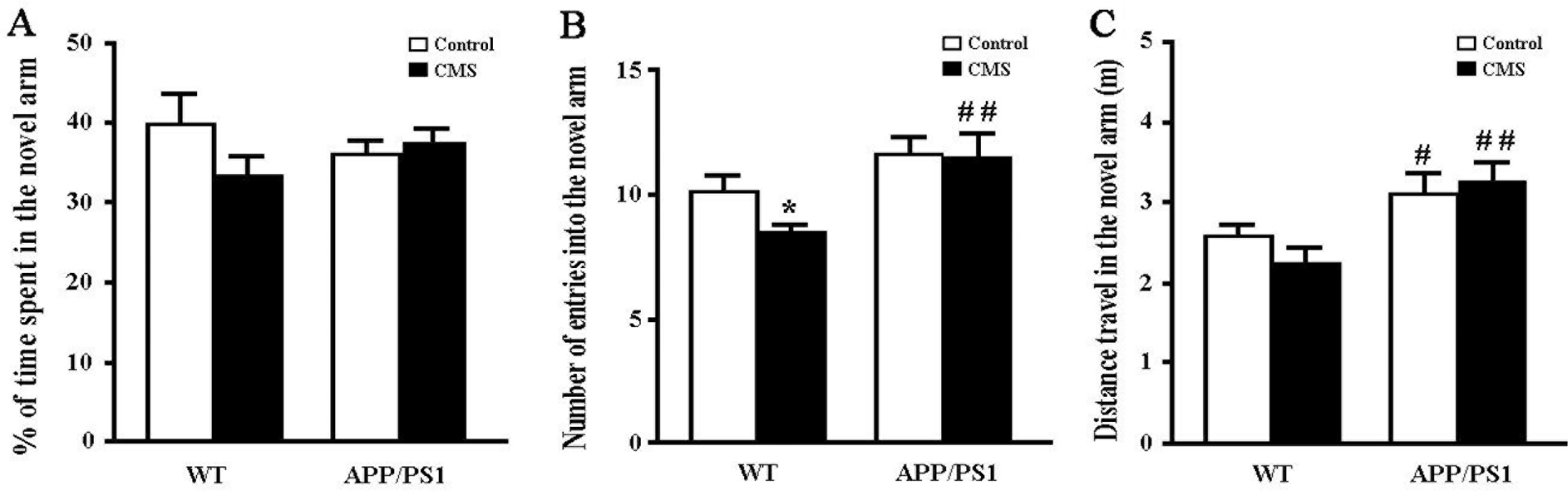
Y maze test of WT and APPswe/PS1dE9 (APP/PS1) mice by CMS for 8 weeks. (A) Percent of time spent in the novel arm (NA). (B) The number of entries into the NA. (C) Total movement distance. Data represent means ± SEM. Two-way ANOVA followed by post-hoc multiple comparison test. *P < 0.05, compared to Control; #P < 0.05; ###P < 0.001, compared to WT. n = 14 in each group.

### The serum CORT concentration increased to the normal levels in APPswe/PS1dE9 mice following CMS exposure

To elucidate the molecular basis of these behavioral changes, we first examined the serum CORT levels of all groups. CMS exposure significantly increased serum CORT concentrations in WT mice, as well as APPswe/PS1dE9 mice (CMS, F_1,20_ = 16.533, P = 0.001; genotype, F_1,20_ = 9.079, P = 0.008; interaction, F_1,20_ = 0.106, P = 0.748). Notably, when compared with WT-Control mice, APPswe/PS1dE9-Control mice naturally exhibited low levels of serum CORT (P < 0.01). Serum CORT increased in APPswe/PS1dE9-CMS mice, but was still less than that in WT-CMS littermates (P < 0.05) (Fig. 5). These data indicates that the despair-like phenotype in APPswe/PS1dE9 mice at the baseline might be not associated with activation of the HPA axis.

**Fig. 5.**
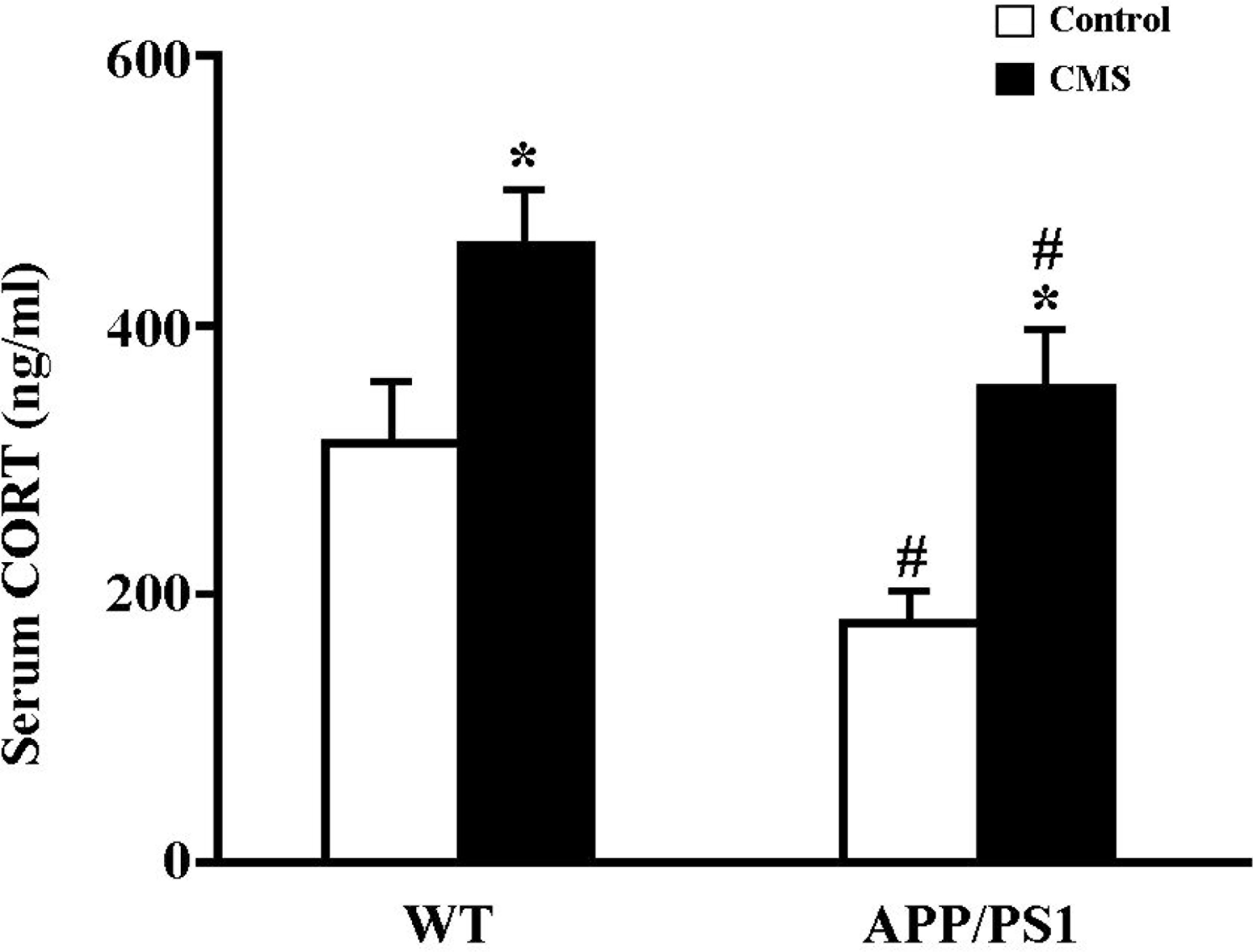
Analysis of serum CORT concentration of WT and APPswe/PS1dE9 (APP/PS1) mice following CMS for 8 weeks. Data represent means ± SEM. Two-way ANOVA followed by post-hoc multiple comparison test. *P < 0.05, compared to Control; #P < 0.05, compared to WT. n = 7 in each group.

### CMS did not exacerbate Aβ accumulation in the hippocampus of APPswe/PS1dE9 mice

Previous studies confirmed that deposition of Aβ plaques in the hippocampus of APPswe/PS1dE9 mice is age-dependent (Trinchese et al., 2004; Hamilton and Holscher, 2012; Strekalova et al., 2015) and aggravated under various pathological conditions, such as hypercholesterolemia (Oksman et al., 2006; Fernández et al., 2009) and transient ischemia (Kemppainen et al., 2014). However, the consequence of CMS on Aβ load and metabolism is still unknown. In order to further explain the significance, we first detected extracellular Aβ deposition in each group. No thioflavin-S-positive fibrillary plaques were present in the hippocampus of APPswe/PS1dE9 mice (Fig. 6A). Previous studies also reported that deposition of diffuse plaques may occur in the brain prior to fibrillary plaques (Rak et al., 2007). To this end, we further performed immunohistochemical staining for 6E10 antibodies that can detect APP and different forms of Aβ peptide fragments (Obregon et al., 2012). No 6E10-immunopositive diffuse plaques were observed in the hippocampus of any group. However, 6E10-immunoreactive product was present in the pyramidal layers and granule layers of APPswe/PS1dE9 mice, but not aggravated by CMS exposure (P > 0.05) (Fig. 6B, C).

**Fig. 6.**
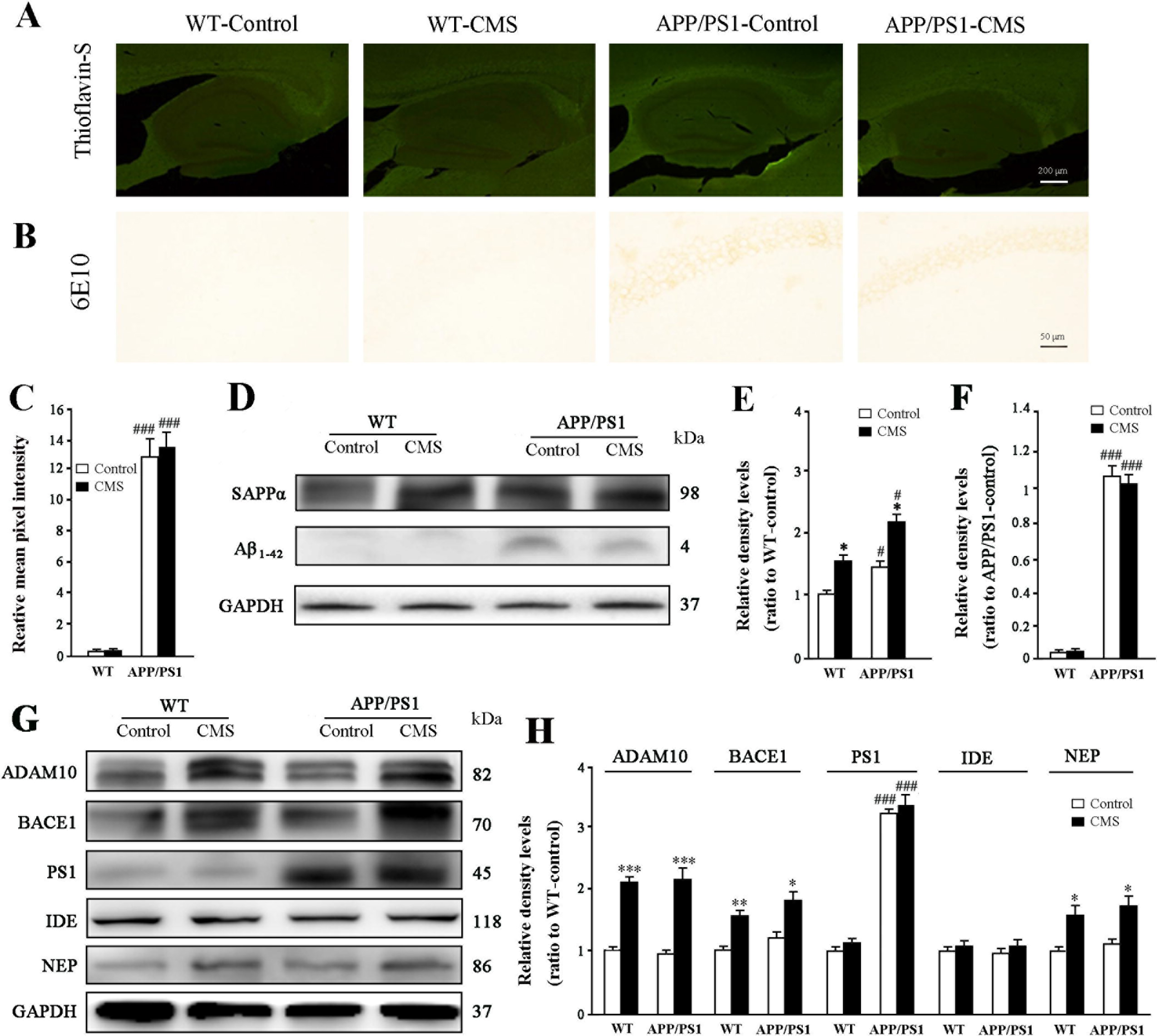
Analysis of Aβ accmulation and metabolism in the hippocampus of WT and APPswe/PS1dE9 (APP/PS1) mice by CMS for 8 weeks. (A) Thioflavin-S staining. (B) 6-E10 immunohistochemical staining. Neither Thioflavin-S-positive fibrillary plaques nor 6E10-immunopositive diffuse plaques were observed in the hippocampus of the both groups of mice. (C) Representative immunoblot and (D-E) corresponding densitometry analysis for soluble amyloid precursor protein-α peptides (sAPPα) and soluble Aβ peptides. (F) Representative immunoblot and (G) corresponding densitometry analysis for APP secretases, including a-disintegrin and metalloproteinase 10 (ADAM10), (β-site amyloid precursor protein-cleaving enzyme 1 (BACE1) and presenilin1 (PS1), and Aβ-degrading enzymes including neprilysin (NEP) and insulin-degrading enzyme (IDE). Data represent means ± SEM. Two-way ANOVA followed by post-hoc multiple comparison test. *P < 0.05; **P < 0.01; ***P < 0.001, compared to Control; #P < 0.05; ###P < 0.001 compared to WT. n = 4 in each group.

Subsequently, we analyzed the effect of CMS on contents of soluble APP α fragment (sAPPα) and soluble Aβ_1-42_ in the hippocampus of WT and APPswe/PS1dE9 mice. Western blot and densitometry analysis revealed that CMS increased expression levels of sAPPα in the both genotypes (both P < 0.05) (CMS, F_1,12_ = 119.319, P < 0.001; genotype, F_1,12_ = 14.259, P = 0.005; interaction, F_1,12_ = 11.576, P < 0.001). APPswe/PS1dE9-Control mice and APPswe/PS1dE9-CMS mice showed higher sAPPα content than WT-Control mice and WT-CMS mice, respectively (both P < 0.01) (Fig 6D, E). The soluble Aβ_1-42_ levels were much higher in APPswe/PS1dE9 mice than WT mice, and not increased following CMS (CMS, F_1,12_ = 0.114, P = 0.745; genotype, F_1,12_ = 48.798, P < 0.001; interaction, F_1,12_ = 0.141, P = 0.717) (Fig 6D, F).

We further analyzed APP secretases, including α-disintegrin and metalloproteinase 10 (ADAM10; α-secretase), β-site amyloid precursor protein-cleaving enzyme 1 (BACE1; β-secretase) and PS1 (γ-secretase) in the hippocampus of different group animals. CMS significantly increased expression levels of ADAM10 and BACE1 (F_1,12_ = 134.997, P < 0.001; F_1,12_ = 55.190, P < 0.001, respectively), but effects of genotype (F_1,12_ = 0.884, P = 0.375; F_1,12_ = 2.514, P = 0.152, respectively) and CMS × genotype (F_1,12_ = 1.180, P = 0.309; F_1,12_ = 0.267, P = 0.619, respectively) were not apparent. As expect, the expression levels of PS 1 were much high in the hippocampus of APPswe/PS1dE9 mice, compared to WT mice (F_1,12_ = 239.739, P < 0.001). But effects of CMS and CMS × genotype were not significant (F_1,12_ = 1.283, P = 0.290; F_1,12_ = 0.004, P = 0.950, respectively) (Fig. 6G, H).

In addition, we analyzed effects of CMS on Aβ degrading enzymes levels, including insulin degrading enzyme (IDE) and neprilysin (NEP), in the hippocampus of WT and APPswe/PS1dE9 mice. IDE expression was not significantly affected by treatment (F_1,12_ = 1.795, P = 0.217), genotype (F_1,12_ = 0.558, P = 0.476) or their interaction (F_1,12_ = 0.607, P = 0.803). However, CMS significantly increased NEP levels in both genotypes of mice (treatment, F_1,12_ = 19.908, P < 0.001; genotype, F_1,12_ = 1.162, P = 0.312; interaction, F_1,12_ = 1.606, P = 0.241) (Fig. 6G, H).

Taken together, APPswe/PS1dE9 mice had elevated levels of sAPPα and Aβ_1-42_ in the hippocampus, which was due to overexpression of PS1, an intramembrane-cleaving protease responsible for the final proteolytic event in the amyloidogenic and nonamyloidogenic pathways (Huang and Mucke, 2012). CMS increased expression levels of BACE1 and NEP, which would maintain the homeostasis between Aβ production and degradation, thereby resulting in no changes in brain Aβ levels of APPswe/PS1dE9 mice.

### CMS increased neuroinflammation in the hippocampus of APPswe/PS1dE9 mice

Neuroinflammation plays a key role in the pathogenesis of AD and depression (Choi and Ryter, 2014). We determined effects of CMS on several inflammatory factors, including IL-1β, IL-6 and TNF-α, in the hippocampus of WT and APPswe/PS1dE9 mice. Results revealed a significant effect of CMS on the expression levels of IL-1β (F_1,12_ = 35.643, P < 0.001), IL-6 (F_1,12_ = 178.172, P < 0.001) and TNF-α (F_1,12_ = 61.174, P < 0.001). Genotype significantly affected IL-6 (F_1,12_ = 57.276, P < 0.001) and TNF-α (F_1,12_ = 23.064, P = 0.001), but did not significantly affect IL-1β (F_1,12_ = 4.727, P = 0.061). CMS × genotype affected IL-1β (F_1,12_ = 83.484, P < 0.001) and TNF-α (F_1,12_ = 14.581, P = 0.005), but not IL-6 (F_1,12_ = 2.373, P = 0.165), significantly. Further post-hoc multiple comparison tests revealed that CMS increased IL-1β, IL-6 and TNF-α in WT mice, but only increased IL-6 and TNF-α in APPswe/PS1dE9 mice (All P < 0.05). Only IL-1β and IL-6 increased in APPswe/PS1dE9-Control mice, when compared with those in WT-Control mice (both P < 0.05) (Fig. 7A, B).

**Fig. 7.**
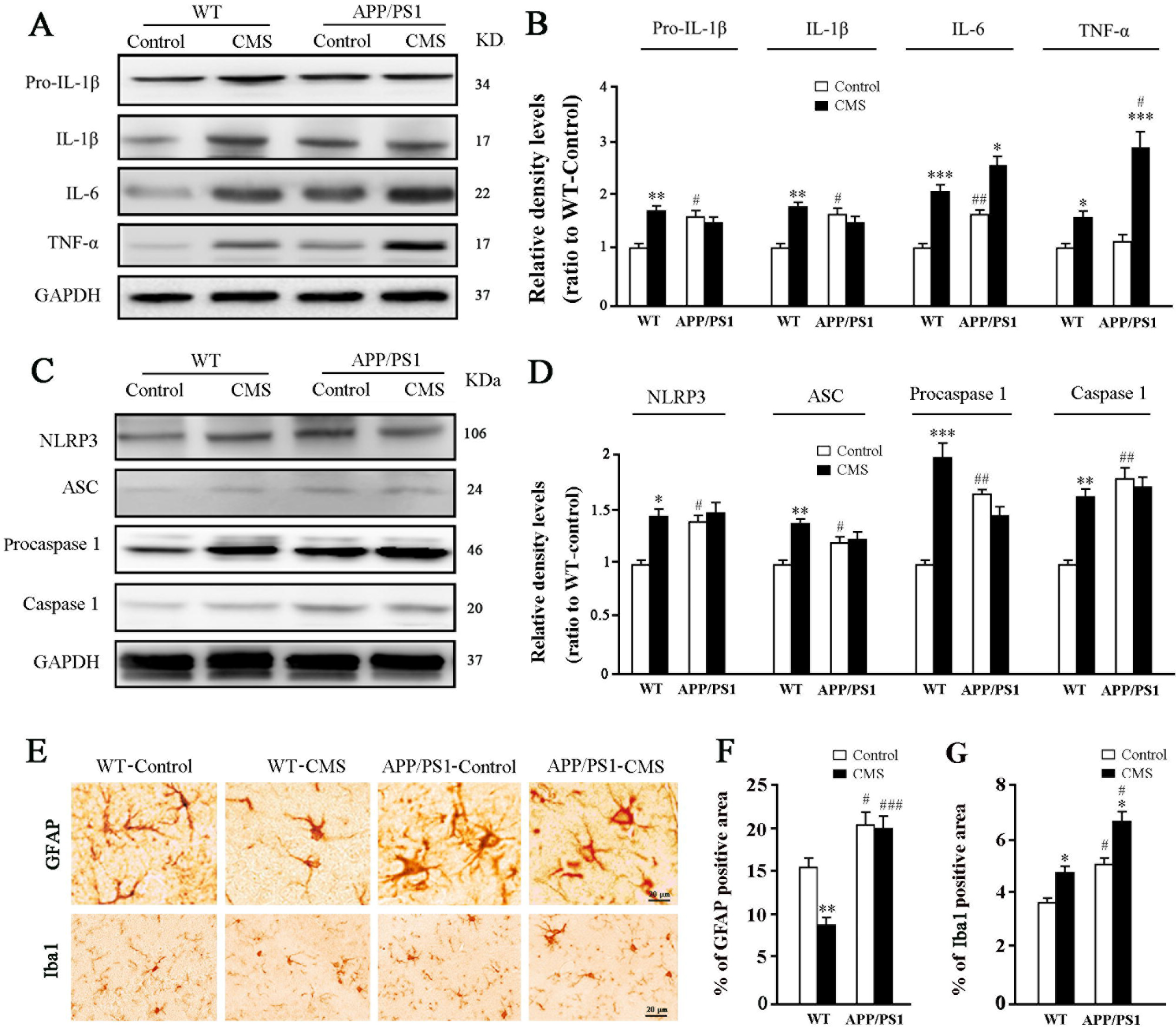
Analyses of neuroinflammation and NLRP3 inflammsome activation in the hippocampus of WT and APPswe/PS1dE9 (APP/PS1) mice following CMS for 8 weeks. (A) Representative immunoblot and (B) corresponding densitometry analysis for pro-interleukin-1β (IL-1β), interleukin-1β (IL-1β), interleukin-6 (IL-6) and tumor necrosis factor-α (TNF-α). (C) Representative immunoblot and (D) corresponding densitometry analysis of proteins involved in the pathway of NLRP3 inflammsome activation. Data represent means ± SEM. Two-way ANOVA followed by post-hoc multiple comparison test. *P < 0.05;**P < 0.01, ***P < 0.001, compared to Control; #P < 0.05; ##P < 0.01; ###P < 0.001, compared to WT. n = 4 for each group.

### NLRP3 inflammasome activation in the hippocampus of APPswe/PS1dE9 mice

Inflammatory factor production is dependent on inflammasome activation; NOD-like receptor containing a pyrin domain 3 (NLRP3) inflammasome mediates neuroinflammation in depression, as well as in AD (Pan et al., 2014; Zhang et al., 2015; Du et al., 2016). WT-CMS mice, APPswe/PS1dE9-Control mice and APPswe/PS1dE9-CMS mice showed high NLRP3 inflammasome activation, as revealed by upregulated expression of NLRP3, apoptosis-associated speck-like protein containing a caspase recruitment domain (ASC), Caspase1 and Procaspase1 (NLRP3: CMS, F_1,12_ = 7.194, P = 0.0028; genotype, F_1,12_ = 4.804, P = 0.060; interaction, F_1,12_ = 4.146, P = 0.076; ASC, F_1,12_ = 17.754, P = 0.003; F_1,12_ = 0.221, P = 0.651; interaction, F_1,12_ = 16.163, P = 0.004; Caspase1: CMS, F_1,12_ = 20.122, P = 0.002; genotype, F_1,12_ = 44.900, P < 0.001; interaction, F_1,12_ = 26.579, P < 0.001; Procaspase1: F_1,12_ = 0.090, P = 0.772; genotype, F_1,12_ = 0.689, P = 0.430; interaction, F_1,12_ = 4.773, P = 0.060) (Fig. 7C, D).

### CMS increased microglial activation in the hippocampus of APPswe/PS1dE9 mice

Activated glia cells are a main source of neuroinflammatory factors in the CNS (Santos et al., 2016), and increased levels of microglia-derived cytokines correlate with AD and depression (Santos et al., 2016). We further addressed alterations of astrocytes and microglia in APPswe/PS1dE9 mice after CMS exposure. Semi-quantitative immunohistochemistry demonstrated that CMS reduced the percentage area of GFAP positive signal in WT mice, indicating the occurrence of astrocyte atrophy (Cobb et al., 2016), but not in APPswe/PS1dE9 mice (CMS, F_1,12_ = 9.188, P = 0.016; genotype, F_1,12_ = 74.441, P < 0.001; interaction, F_1,12_ = 5.826, P = 0.042). APPswe/PS1dE9-CMS mice had high percentage area of glial fibrillary acidic protein (GFAP) expression compared to APPswe/PS1dE9-Control mice (P < 0.05), but the difference was not significant between APPswe/PS1dE9-CMS and APPswe/PS1dE9-control mice (P > 0.05) (Fig 7E, F). CMS caused microglial activation in both WT and APPswe/PS1dE9 mice, revealed by increased percentage area of ionized calcium-binding adaptor molecule 1 (Iba1) expression (CMS, F_1,12_ = 13.396, P = 0.006; genotype, F_1,12_ = 15.242, P = 0.005; interaction, F_1,12_ = 0.259, P = 0.625) (Fig. 7E, F).

### Downregulation of BDNF-TrkB-CREB signaling pathway in the hippocampus of APPswe/PS1dE9 mice

Downregulation of BDNF-TrkB-CREB (cAMP response-element binding protein) signaling pathway is one of the main pathophysiological changes in depression and AD (Berry et al., 2012; Jeanneteau et al., 2012). We demonstrated that levels of BDNF, TrkB and p-CREB were significantly decreased by CMS (F_1,12_ = 26.757, P < 0.001; F_1,12_ = 61.263, P < 0.001, F_1,12_ = 44.369, P < 0.001, respectively), overexpression of transgenic APPswe/PS1dE9 (F_1,12_ = 10.998, P = 0.011; F_1,12_ = 39.145, P < 0.001, F_1,12_ = 45.823, P < 0.001, respectively) and their interaction (F_1,12_ = 10.270, P = 0.013, F_1,12_ = 32.069, P < 0.001, F_1,12_= 39.357, P < 0.001, respectively). The expression levels of BDNF, TrkB and p-CREB were not different among WT-CMS, APPswe/PS1dE9-Control and APPswe/PS1dE9-CMS mice, but were less than those in WT-Control mice (all P < 0.05) (Fig. 8A, B).

**Fig. 8.**
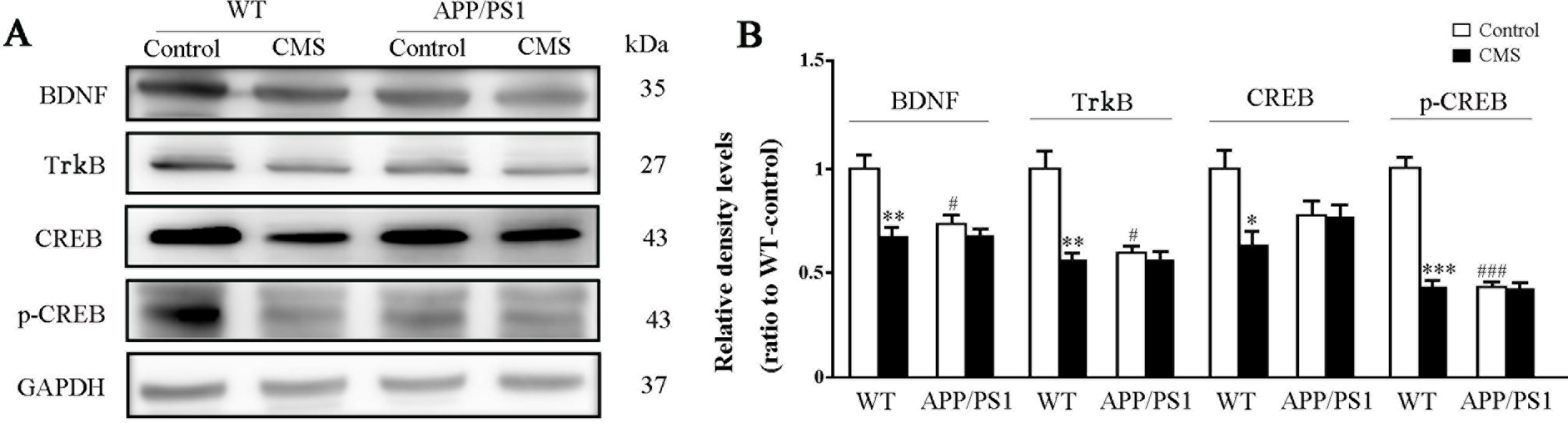
Activation of BDNF signaling pathway in the hippocampus of WT and APPswe/PS1dE9 (APP/PS1) mice by CMS for 8 weeks. (A) Representative immunoblot and (B) corresponding densitometry analysis 47, for BDNF, TrkB, CREB and p-CREB. Data represent means ± SEM. Two-way ANOVA followed by post-hoc multiple comparison test. *P < 0.05;**P < 0.01; ***P < 0.001, compared to Control; #P < 0.05; ###P < 0.001, compared to WT. n = 4 for each group.

**Fig. 9.**
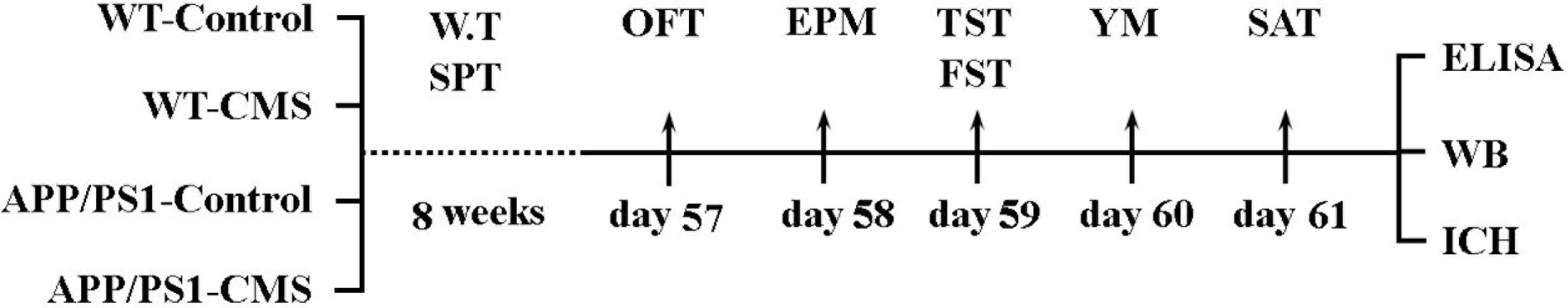
Diagram depicts animal grouping and experimental procedures. Eight-week old WT and APPswe/PS1dE9 (APP/PS1) mice received CMS or no treatment for 8 weeks, during which the weight test (W.T) and sucrose preference test (SPT) were performed per week. From 57^th^ day to 61^st^ day, serial behavioral tests including the open field test (OFT), elevated plus maze (EPM), tail suspension test (TST), forced swimming test (FST), Y-maze and social approach test (SAT) were carried out in the sequence. The mice were then sacrificed on the next day after the last behavioral test, followed by ELISA, immunohistochemistry (IHC) and Western blot (WB) analyses.

## DISCUSSION

There has been no effective cure during the progressive stage of AD. Thus, early diagnosis of AD is an attractive strategy to increase the survival rate of the patients. To achieve this goal, it is necessary to identify subtle abnormal symptoms and related pathophysiological changes during the early stages of AD. The present study demonstrated that at baseline, 4-month-old APPswe/PS1dE9 mice exhibit a series of non-cognitive symptoms, such as behavioral despair, social disability, anxiety and hyperactivity when they are yet to display cognitive dysfunctions, thus better simulating the clinical features of patients with the early stage AD. Furthermore, we found that, in comparison with WT-Control mice, both WT-CMS mice and APPswe/PS1dE9 mice demonstrate neuroinflammation and neurotrophic disturbance, potentially serving as common pathological mechanisms between AD and depression. Interestingly, CMS does not significantly increase behavioral despair in APPswe/PS1dE9 mice, but increases anxiety-like behaviors, and its pathological basis might be related to increased neuroinflammation. These findings aid in understanding the pathophysiological mechanisms of mental symptoms during the initial stages of AD, which would contribute to the early diagnosis of the disease.

APPswe/PS1dE9 mice are one of the most commonly used AD model animals for investigating Aβ-related pathogenesis and preclinical efficacy evaluation against AD. Previous studies on the behaviors of APPswe/PS1dE9 mice mainly focus on the aspects of cognitive dysfunction. Only limited literature is available on the non-cognitive abnormalities of these AD model mice (Pietropaolo et al., 2012; Bilkei-Gorzo et al., 2014; Janus et al., 2015; Verma et al., 2015; Olesen et al., 2016). For example, Filali and colleagues (2011) reported that 6-month-old APPswe/PS1dE9 mice exhibit reduced social interaction and hyperactivity. Our previous study reported that 17-month-old APPswe/PS1dE9 mice placed in isolation housing for 3 months exhibit more severe social withdraw (Huang et al., 2015). In the present study, we completed a systematic phenotypic analysis of behavioral and psychological symptoms of this AD model before deposition of Aß plaques. Compared with their age-match WT littermates, 4-month-old APPswe/PS1dE9 mice show increased duration of immobility in the TST and FST, which reflects more severe despair. Our results further revealed that impaired social interaction occurs prior to cognitive dysfunction. We also confirmed that young APPswe/PS1dE9 mice have decreased exploration in the center area of the OFT and the open arm of elevated plus maze, which are widely accepted as the anxiety-like behaviors of mice. In addition, APPswe/PS1dE9 mice exhibit hyperactivity, which is similar to the increased wandering behavior in AD patients (Rolland et al., 2005). Together, these behavioral data highlight that young APPswe/PS1dE9 mice partially mimic depression-like and anxiety-like symptoms in patient with early AD.

There is a lack of some literature investigating specific pathological mechanisms of psychological behaviors of AD genetic mouse models. Our results reveal that 4-month-old APPswe/PS1dE9 mice and WT-CMS mice share similar depression-like behaviors. However, unlike WT-CMS mice, young APPswe/PS1dE9 mice show normal preference for sucrose and body weight gain. Moreover, the behavioral despair of APPswe/PS1dE9 mice seems not to be associated with overactivation of the HPA axis. Contrary to significantly high CORT levels in the blood of WT-CMS mice, 4-month-old APPswe/PS1dE9-Control mice have low serum CORT compare to WT-Control ones. After CMS, serum CORT is increased and returned back to the normal level. This is consistent with the clinical study demonstrating that AD patients and elderly depressive patients show different serum dexamethasone levels and respond differently to the dexamethasone suppression test (Molchan et al., 1990).

AD and major depressive disorder share similar pathologies in the hippocampus and prefrontal cortex. Apart from Aβ and tau load, neurotrophic factor deficiency, impaired neurogenesis, and neuroinflammation are present in AD (Tanzi and Bertram, 2005; Huang and Mucke, 2012), all of which are involved in major depressive disorder (Zhang et al., 2016). For example, increased TNF-α and IL-6 levels in peripheral blood correlate robustly with major depressive disorder (DowlatiSantos et al., 2010) and AD (Swardfager et al., 2010). Furthermore, AD patients with comorbid depression show high levels of circulating IL-6 and TNF-α, compared to non-depressed AD patients (Khemka et al., 2014).

Changes in BDNF signaling has been implicated both in the etiology of depression and in antidepressant drug action (Castrén and Rantamäki, 2010). Consistently, higher cortisol levels have been shown to lower BDNF in the hippocampus of rats, with antidepressants reversing the change (Haynes, et al., 2004). Reduced levels of BDNF the hippocampus and cerebral cortex are also well documented in AD, even during the pre-clinical stages (Peng et al., 2005). In agreement with this view, decreases in BDNF levels, and increases in IL-1β and IL-6 levels with NLRP3 inflammasome activation, are observed in the hippocampus of both APPswe/PS1dE9-Control mice and WT-CMS mice. Therefore, overexpression of transgenic APP/PS1 causes high levels of Aβ, subsequently results in dysregulation of BDNF-signal pathway and neuroinflammation, which may be the main pathogenesis for non-cognitive psychological behaviors of this AD mouse model.

Depression and anxiety share high rates of co-morbidity, with both incorporating an underlying element of stress (EI Yacoubi et al., 2013; Farrell and O’Keane, 2016). However, different molecular mechanisms between depression and anxiety are not clear. Chronic injection of CORT specially induces depression-like behaviors, which can be prevented by mineralocorticoid receptor antagonist (Wu et al., 2013). A recent study revealed that the absence of CORT production by adrenalectomy completely cancels depression-like behaviors, but does not normalize anxiety-like behaviors after CMS exposure (Chen et al., 2016). The present study provides additional experimental evidences for separating anxiety from behaviors related to depression. CMS aggravates anxiety-like behaviors, but not depression-like behaviors in the early stages of APPswe/PS1dE9 mice that exhibit low serum CORT levels at baseline condition. The biochemical and molecular analyses revealed that CMS normalizes CORT levels and increasesIl-6 and TNF-α, but does not affect BDNF and Aβ levels in the hippocampus of APPswe/PS1dE9 mice. These data suggest that increased neuroinflammation may be one of the contributing factors in aggravating anxiety-like symptoms of APPswe/PS1dE9-CMS mice. Indeed, accumulative evidence has highlighted a close relationship between inflammation and mental diseases (Zunszain et al., 2013; Santos et al., 2016; Wohleb et al., 2016; Bialas et al., 2017). Further study is necessary to identify the exact effect of anti-inflammatory therapy on the alleviation of mental symptoms in early AD.

In summary, this study has characterized the non-congitive symptoms as well as several cellular, molecular and neuropathological alterations in APP/PS1 mice before amyloidosis onset. Furthermore, the results suggest that neuroinflammation and BDNF disturbance are common pathological mechanisms of depression/anxiety behaviors in the CMS model and APPswe/PS1dE9-Control mice, but the causes are different. The former may be due to stress induced-activation of the HPA axis, and the latter due to the increase of Aβ in the brain. This finding provides an experimental pathological basis for distinguishing the mental symptoms with early AD from major depression.

## MATERIALS AND METHODS

### Animal grouping

Three months old male APPswe/PS1dE9 transgenic mice with a C57BL/6J background were obtained from the Model Animal Research Center of Nanjing University. These transgenic mice were crossed with C57BL female mice to produce the offspring APPswe/PS1dE9 mice and their wild-type (WT) littermates. The mouse genotype was identified using polymerase chain reaction, as previously (Jankowsky et al., 2001). After weaning, mice were housed in same-sex groups of 3-4 under standard laboratory conditions (12-hour light-dark cycle) with a room temperature of 22 °C, and water and food available ad libitum.

At 8 weeks old, the offspring male and female APPswe/PS1dE9 mice and their WT littermates were randomly divided into four groups: WT-Control (n = 14), WT-CMS (n = 20), APPswe/PS1dE9-Control (n = 14) and APPswe/PS1dE9-CMS (n = 20). Animal experiments were performed in accordance with the Guide for the Care and Use of Laboratory Animals of Nanjing Medical University. Fig. 1 presents the experimental schedule and timeline for the study.

### CMS

Control mice were housed under standard conditions, whereas stressed animals were exposed to the CMS procedure for a period of 56 consecutive days, which has been established well in our laboratory (Kong et al., 2008; Du et al., 2016). The CMS regimen consisted of once or twice daily exposure to different stressors including inversion of day/night light cycle, soiled cage bedding, 45° tilted cage, overnight food and water deprivation or restraint and so on. The sucrose preference and body weight of each mouse were measured weekly during this period.

### Sucrose preference test

The sucrose preference test was used to detect anhedonia of depressive animals (Du et al., 2016). Mice were given the choice to drink from 2 different bottles for 24 h: one contained a sucrose solution (1%) and the other contained only tap water. The positions of the bottles in the cage were placed randomly and switched every 8 h. The preference for sucrose was measured as a percentage of the consumed sucrose solution relative to the total amount of liquid intake.

### Forced swim test

Mice were forced to individually swim in an open cylindrical container (diameter: 15 cm; height: 25 cm) with 14 cm of water at room temperature (about 22 ± 1°C) for 6 min (Du et al., 2016). The animals were considered immobile when they did not attempt an escape, but instead floated upright with only small movements necessary to keep their head above water. The last 4 min was scored for the immobility time by Tail Susp Scan (Clever Sys).

### Tail suspension test

Mouse tails were wrapped with tape from the base to the tip, covering about four-fifths of their length and fixed upside down on the hook for 6 min (Du et al., 2016). The mice were considered immobile when they hung passively and completely motionless. The duration of immobility was observed and measured during the last 4 min Tail Susp Scan (Clever Sys).

### Social sociability test

The apparatus of social sociability testing included three plastic compartments (69 cm × 13 cm × 15 cm), separated by two doors (8 cm × 8 cm) (Huang et al., 2015). Each experimental mouse was placed into the central compartment and allowed to explore the apparatus for a 5-min acclimatization period. Then, an unfamiliar adolescent C57BL female mouse (stranger-1), was introduced into the circular wire mesh cage (diameter: 9 cm, height: 15 cm) of either lateral compartment. The tested mouse was returned to the central compartment and explored all three compartments for another 5 min. The time spent in each compartment was collected, and data were expressed as the ratio between time spent with an unfamiliar mouse and time in the empty enclosure.

### Open field test

The open field test was used to evaluate anxiety-like behaviors (Huang et al., 2015). The open field consisted of a square black Plexiglas box (60 cm × 60 cm × 25 cm), with an outlined center area (30 cm × 30 cm). Each animal was placed in the middle of box, serving as a starting point, then allowed to move freely for 5 min within the box. The amount of time and distance traveled in the center area of the maze were measured.

### Elevated plus maze test

The elevated plus maze, made of two open arms (50 cm × 10 cm) and two closed arms (50 cm × 10 cm × 40 cm), was used to test the anxiety of animals (Huang et al., 2016). For this experiment, each mouse was first placed into the closed arms and allowed to explore the maze freely for 5 min. The percentage of entries into, time spent and distance travelled in the open arms, were recorded.

### Y-maze test

The Y-maze test was performed to measure short-term spatial memory of mice (Huang et al., 2016). The Y-maze consists of three arms (40 cm long, 12 cm high, 3 cm wide at the bottom and 10 cm wide at the top) with an angle of 120° between the arms. The test includes two 5-min stages, with an interval of 2 h. During the first stage, one arm, considered as the novel arm, was blocked, allowing mice to randomly explore the two other arms. During the second stage, the novel arm was open, and the mice could freely explore all three arms. The number of entries into and percentage of time traveled in each arm were recorded. The travelling speed was also calculated.

Mouse activity in the aforementioned behavioral apparatuses was collected by a digital video camera connected to a computer-controlled system (Beijing Sunny Instruments Co. Ltd, China). All behavioral tests were performed by two independent experimenters blind to the identity of experimental groups.

### Mouse tissue preparation

On the next morning after behavioral tests completed, mice were anesthetized. Blood samples of half of each group were collected by eyeball removal for ELISA assay of CORT levels. Subsequently, those mice were sacrificed by cervical dislocation and brains were removed for Western blotting analyses. The remaining mice were transcardially perfused with 0.9% saline by perfusion pump (Cole-parmer) for 5 min, followed by 4% paraformaldehyde for 12 min. The brains were removed, post-fixed 2 h at 4 °C, then dehydrated in a series of graded ethanol solutions, and finally embedded in paraffin. Serial sections were cut sagittally at 5 μm using a paraffin slicing machine (LeicaRM2135, Nussloch, Germany) for pathological analysis.

### ELISA analysis of serum CORT

Blood samples were allowed to clot for 4 h at room temperature, after which the samples were centrifuged for 10 min at 3000 rpm. The serum was obtained and subpackaged (50 μL per tube) and stored at −80 °C until use. Serum CORT levels were detected by ELISA kits (Wuhan Cloud Clone Corp, Hubei, China). The test was performed following the protocol recommended by the manufacturer. Serum concentrations of CORT were calculated using a linear equation, derived from log it transformation of the absorbance and log concentrations of the standards.

### Immunohistochemistry

Immunohistochemical staining was conducted as previously described (Huang et al., 2016). Briefly, after deparaffinization and rehydration, sections were incubated with one of the following primary antibody: mouse monoclonal 6E10 antibody against amino acids 1–17 of Aβ peptide (1:1000; Covance), mouse monoclonal antibody against glial fibrillary acidic protein (GFAP) (1:800; Millipore) or rabbit polyclonal antibody against ionized calcium-binding adaptor molecule 1 (Iba1) (1:1000; Wako), at 4 °C overnight. Following by washing with PBS, sections were then incubated with biotinylated-conjugated goat anti-mouse or rabbit IgG (1:200, Vector Laboratories) for 1 h at 37 °C. Results were visually evaluated after staining with diaminobenzidine (Sigma-Aldrich, USA). The micrographs were captured by a digital microscope system (Leica Microsystems, Wetzlar, Germany).

### Imaging analysis

GFAP, Iba1 or 6E10 immunostained hippocampal sections were photographed at 200× magnification using a digital microscope (Leica Microsystems) The percentage of GFAP or Iba1-immunopositive area was measured using NIH Image J software as described previously (Xu et al., 2015). Briefly, the hippocampal area in each section was manually delineated. The area of positive signal was measured using the interest grayscale threshold analysis with constant settings for minimum and maximum intensities for each staining marker. The percentage area of positive signal was calculated by dividing the area of positive signal to the total area in the region of interest. The mean pixel intensity (MPI) of 6E10-immunoreactivity within pyramidal layers and granule layers was also measured by Image-Pro Plus Software. The immunohistochemical control sections incubated with nonimmune mouse serum were analyzed first, and their MPI was assigned a base value of 0. The relative MPI of each section stained with antibody against 6E10 was then calculated. Four sections per mouse, and 6 mice per group, were used to provide a mean value for each index mentioned above. All quantification was completed by an experimenter who was blind to the identity of the images.

### Western blotting

For Western blot analyses, homogenized samples were loaded onto SDS gels, and then transferred to PVDF membrane. After being blocked with 5% milk, the bands were incubated with rabbit polyclonal antibody against ADAM10 (1:1000; Millipore), ASC (1:1500; Santa Cruz), BDNF (1:300; Abcam), CREB (1:1000; Cell Signaling Technology), Caspase1 (1:1000; Millipore), IDE (1:800; Abcam), IL-1β (1:1000; Millipore), IL-6 (1:1000; Abcam), NEP (1:800; Millipore), PS1 (1:1000; Sigma), NLRP3 (1:1000; AdipoGen), p-CREB (1:1000; Cell Signaling Technology), procaspase 1 (1:500; Millipore), sAPPα (1:800; IBL), TrkB (1:500; Santa Cruz), TNF-α (1:1000; Abcam), rabbit monoclonal antibody against Aβ_1-42_ (1:1000; Abcam), or mouse monoclonal antibody against BACE1 (1:1000; Millipore) at 4°C overnight. Following TBST washing, bands were incubated with horseradish peroxidase-conjugated goat anti-rabbit IgG (1:2000; Vector Laboratories), and then visualized with ECL plus detection system. Protein loading and transfer efficiency was balanced with β-tubulin or GAPDH.

### Statistical analysis

Data are expressed as mean ± SEM. Differences among experimental groups were determined by two-way ANOVA followed by Newman-Keuls post-hoc multiple comparison test. Differences were considered significant at P < 0.05.

## Competing interests

The authors declare no conflict of interest.

## Author contributions

Conceptualization: M.X.; Methodology: J.Y.G., Y.C.; Formal analysis: D.Y.S.; Investigation: J.Y.G., Y.C., D.Y.S., M.X. Writing-original draft: J.Y.G., Y.C., M.X.; Writing review & editing: C.M., M.X.; Visualization: D.Y.S. C.M.; Supervision: M.X.; Project administration: M.X.; Funding acquisition: M.X.

## Funding

This work was supported by grants from the National Natural Science Foundation of China (81671070 and 81271210) and the Natural Science Foundation of Jiangsu Educational Department (14KJA320001).

